# Chronic short sleep in early life accelerates cognitive decline via disrupted proteostasis

**DOI:** 10.64898/2026.03.26.714554

**Authors:** Robert Komlo, Kamalini Sengupta, Ewa Strus, Nirinjini Naidoo

## Abstract

Chronic short sleep (CSS) is an emerging public health issue that frequently begins in adolescence and is common among healthcare professionals and others engaged in shift work. Epidemiological studies associate CSS and sleep disruption with metabolic disorders, cardiovascular disease, cognitive decline, and heightened Alzheimer’s disease risk. Building on our prior findings that sleep deprivation perturbs proteostasis and activates endoplasmic reticulum (ER) stress pathways, we investigated the long-term consequences of CSS in young adult wild-type mice over the course of one year. Mice exposed to CSS displayed impaired cognition in hippocampal dependent tasks by 28 weeks of age, indicating emerging memory deficits. At the molecular level, CSS disrupted hippocampal proteostasis—particularly protein folding processes—and triggered ER stress and activation of the unfolded protein response (UPR). Importantly, disrupted proteostasis preceded the behavioral decline, with diminution of the key chaperone and UPR regulator BiP occurring at 20-22 weeks of age. CSS also increased markers of cellular stress and neuroinflammation while reducing the expression of proteins associated with memory function. Age also seemed to be a cellular stressor, causing a longitudinal increase in UPR, ISR, and neuroinflammation markers. Together, these results indicate that both chronic short sleep and age compromise proteostasis and promote neuroinflammation, contributing to progressive cognitive dysfunction.

## INTRODUCTION

Chronic short sleep (CSS) is pervasive in modern societies. The Center for Disease Control estimates that about half of all middle school children and almost two-thirds of high school students regularly obtain less than the recommended sleep duration (Wheaton AG et al., 2018). More than 36% of adult workers report getting less than 7 hours of sleep/24 hours (Shockey TM et al., 2017). Several variables combine to make up the sleep environment, including light, noise, and temperature. Contributing factors to the burgeoning problem of sleep insufficiency and poor sleep quality include the increased exposure to light from LED bulbs, digital tablets, TV screens, and phones at night, as well as shift work and other work environmental factors.

Epidemiological studies indicate that short sleep, poor sleep quality, and disrupted sleep are all associated with metabolic dysfunction (Wheaton AG et al., 2018), cardiovascular risk (Nagai et al., 2010), cognitive impairments (Kreutzmann et al., 2015) and increased risk for AD (see (Ju et al., 2014) and references therein). Sleep loss and disruption have been shown to accelerate features of brain aging and Alzheimer’s disease (AD) (Musiek and Holtzman, 2016). Acute sleep deprivation increases brain interstitial fluid (ISF) amyloid-beta (Aβ) concentrations and chronic sleep deprivation accelerates Aβ deposition into insoluble amyloid plaques in two transgenic mouse models of amyloidosis (Rothman et al., 2013; Zhao et al., 2017). Sleep loss and sleep disruption also result in cognitive deficits, such as impaired attention, decision-making, learning, and various types of memory (Kreutzmann et al., 2015). Chronic short sleep, such as that experienced by shift workers, imparts neurodegeneration in select populations of neurons as well as increased amyloid production (Zhang et al., 2014; Zhu et al., 2016; Zhu et al., 2015). The cellular and molecular mechanisms by which sleep loss influences neural injury and neurodegeneration are not clearly known.

We have previously shown that sleep loss and sleep disruption perturb proteostasis especially in the endoplasmic reticulum (ER), leading to the induction of a signal transduction pathway, the unfolded protein response (UPR) (Brown et al., 2014; Naidoo et al., 2008; Naidoo et al., 2005). The UPR is an adaptive response that restores proteostasis by attenuating protein translation, facilitating protein degradation, and increasing protein chaperone activity through the activation of Protein Kinase R-like ER Kinase (PERK), inositol-requiring enzyme 1 (IRE1) and activating transcription factor 6 (ATF6) (Hetz et al., 2020; Walter and Ron, 2011). However, sustained activation of the UPR and especially of PERK leads to the integrated stress response (ISR) and downstream activation of inflammatory and pro-apoptotic pathways (Pakos-Zebrucka et al., 2016). Notably, PERK activation regulates protein synthesis, which is known to be important for memory and cognition (Freeman and Mallucci, 2016). Sustained ER stress has been shown to be involved in memory impairments and alleviating ER stress is known to improve memory (Hafycz et al., 2022; Hafycz et al., 2023; Halliday and Mallucci, 2014; Halliday and Mallucci, 2015). However, the temporal resolution of ER stress induction preceding cognitive impairments is not well understood or known. To investigate the relationship between ER stress and cognition, we examined how chronic short sleep in mice during early adulthood affects cognitive performance over the course of a year. Our findings indicate that chronic short sleep triggers and sustains ER stress, disrupting proteostasis before cognitive deficits emerge, while also accelerating cognitive decline.

## RESULTS

### Chronic short sleep accelerates cognitive decline

We assessed the effect of chronic short sleep during early adulthood on cognitive performance across 1-year of aging. Mice subjected to 8 weeks of chronic short sleep (CSS) for 3 days/week as previously described (Owen et al., 2021) and illustrated in Figs. 1A and 1B were tested for hippocampal dependent learning across aging using a spatial object recognition assay (SOR), illustrated in Fig. 1C. We found that these mice showed a decline in cognitive performance earlier than rested mice that underwent the same SOR tests. Initially, SOR testing at 4- and 8-weeks post CSS revealed no differences between the rested and sleep deprived mice in the place preference test. All groups showed increased preference for the moved object during the testing phase, suggesting that any cognitive effects of CSS were not yet visible. However, during the third testing period at 28 weeks of age the rested mice displayed increased place preference for the moved object during the testing period while the CSS mice did not (See Figure 2). The CSS mice as illustrated by the discrimination index, which was used to quantify learning in the SOR, were not able to discriminate between the moved and unmoved objects (Fig. 2C; T-test, p=0.049) (Naidoo et al., 2018; Sivakumaran et al., 2018). The rested mice exhibited a strong preference for the moved object in the SOR test over aging until mice reached 52 weeks when the rested mice did not display any improvement in cognitive performance during the test when compared to the training period. The performance of the 52-week-old rested mice was not significantly different from that of the CSS mice.

**Fig. 1:**
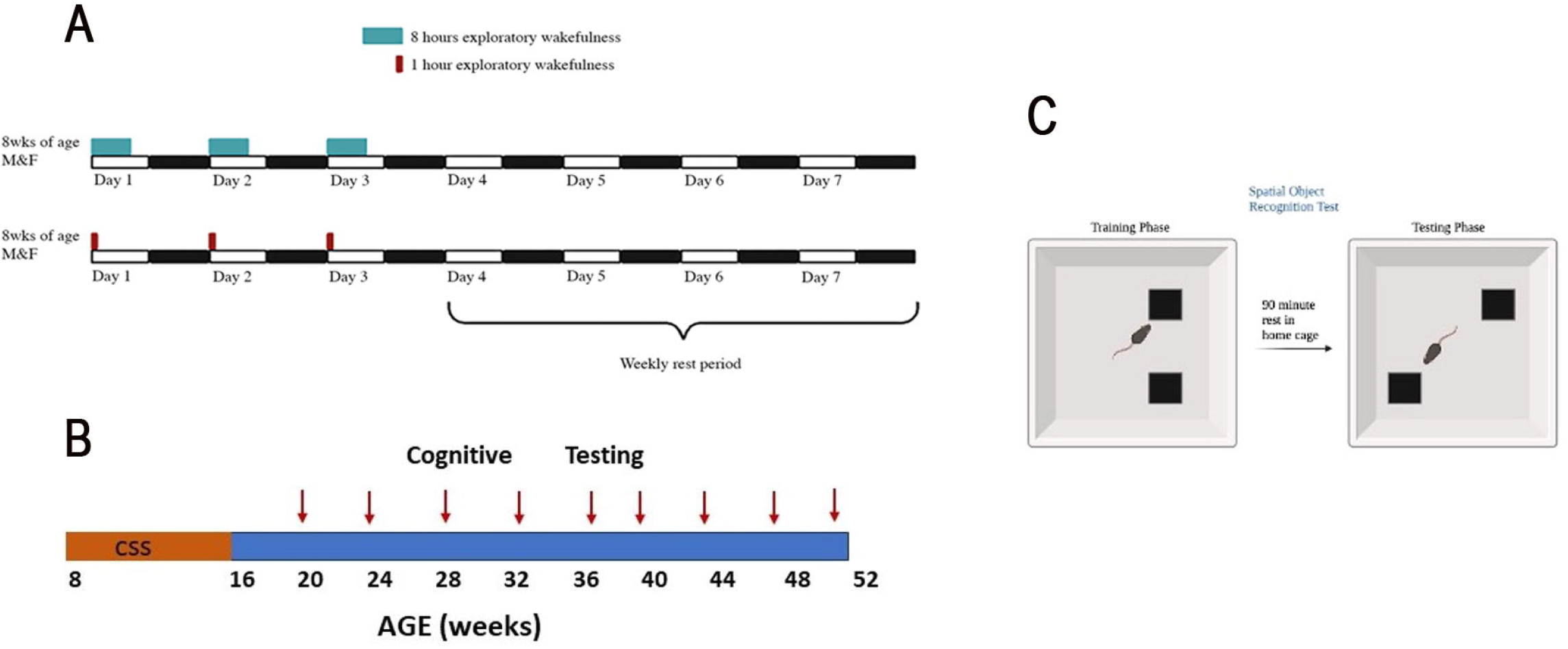
Sleep Deprivation Paradigm used to model CSS. A: Schematic modelling weekly sleep deprivation. CSS mice were sleep deprived for 8 hours 3 out of 7 days of the week, while rested mice were exposed to the same environment for 1 hour 3 out of 7 days of the week. B: Timeline of entire experiment. Arrows indicate the points at which SOR was performed. C: Schematic of SOR test.

**Fig. 2:**
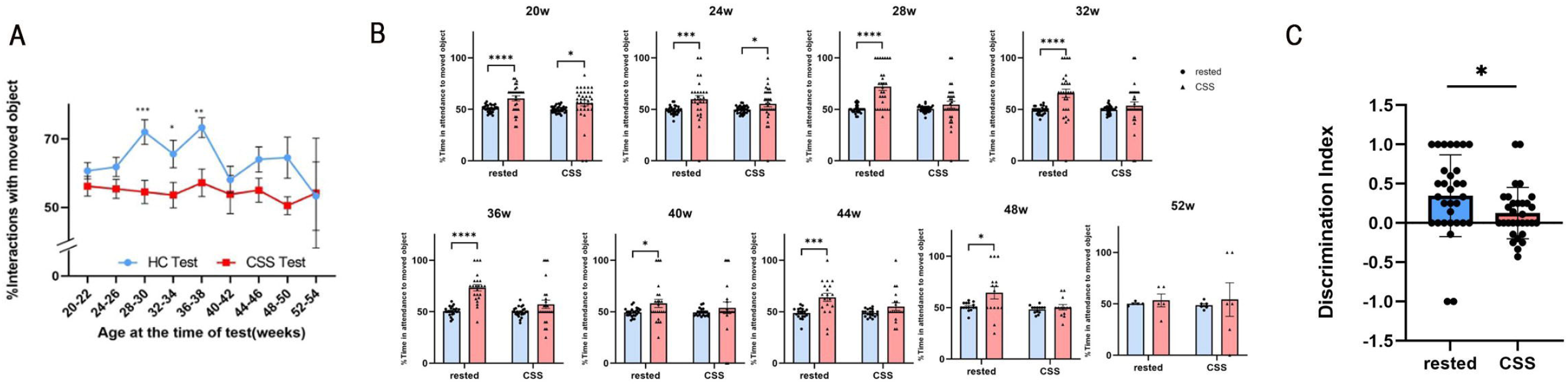
CSS causes cognitive decline through decreased performance in SOR starting at 28 weeks of age. **A:** Temporal profile of cognitive performance of CSS and rested mice. SOR tasks measured using average time in attendance to moved object ± SEM shown for each of the time points, n=20-25, **p<0.01, ***p<0.001. **B:** CSS mice perform worse at SOR started at 28 weeks. Histograms illustrate %time interacting with moved object during training (blue) and during testing (pink). Rested mice show preference for the moved object from 20-48 weeks, while CSS mice only show preference until 24 weeks. Mean and SEM shown for each phase; n=20-25, *p<0.05, ****p<0.0001. C: Discrimination index shown for CSS and rested mice at 28 weeks of age.

### Decline in BiP and impaired proteostasis precedes cognitive impairment

We have previously demonstrated that BiP protein expression declines with age (Naidoo et al., 2008; Naidoo et al., 2018) and that decreased expression of BiP correlates with learning deficits (Hafycz et al., 2022; Hafycz et al., 2023). Based on those observations we examined hippocampal BiP protein expression by immunofluorescence at various timepoints over the one-year experimental period. BiP protein expression is significantly lower in the hippocampi of 20-22 week-old CSS mice when compared to that in rested mice and is also lower at subsequent time points examined (Fig. 3A; unpaired t-test, p=0.010). Importantly, our data indicate that the diminution of BiP precedes the decline in cognitive performance in the CSS mice, which occurs at 28 weeks of age. Our data also indicate that BiP in the rested mice also declines with age (Fig. 3A; 2-way ANOVA, p=0.026), recapitulating previously published data (Hafycz et al., 2022; Naidoo et al., 2008; Naidoo et al., 2018).

**Fig. 3:**
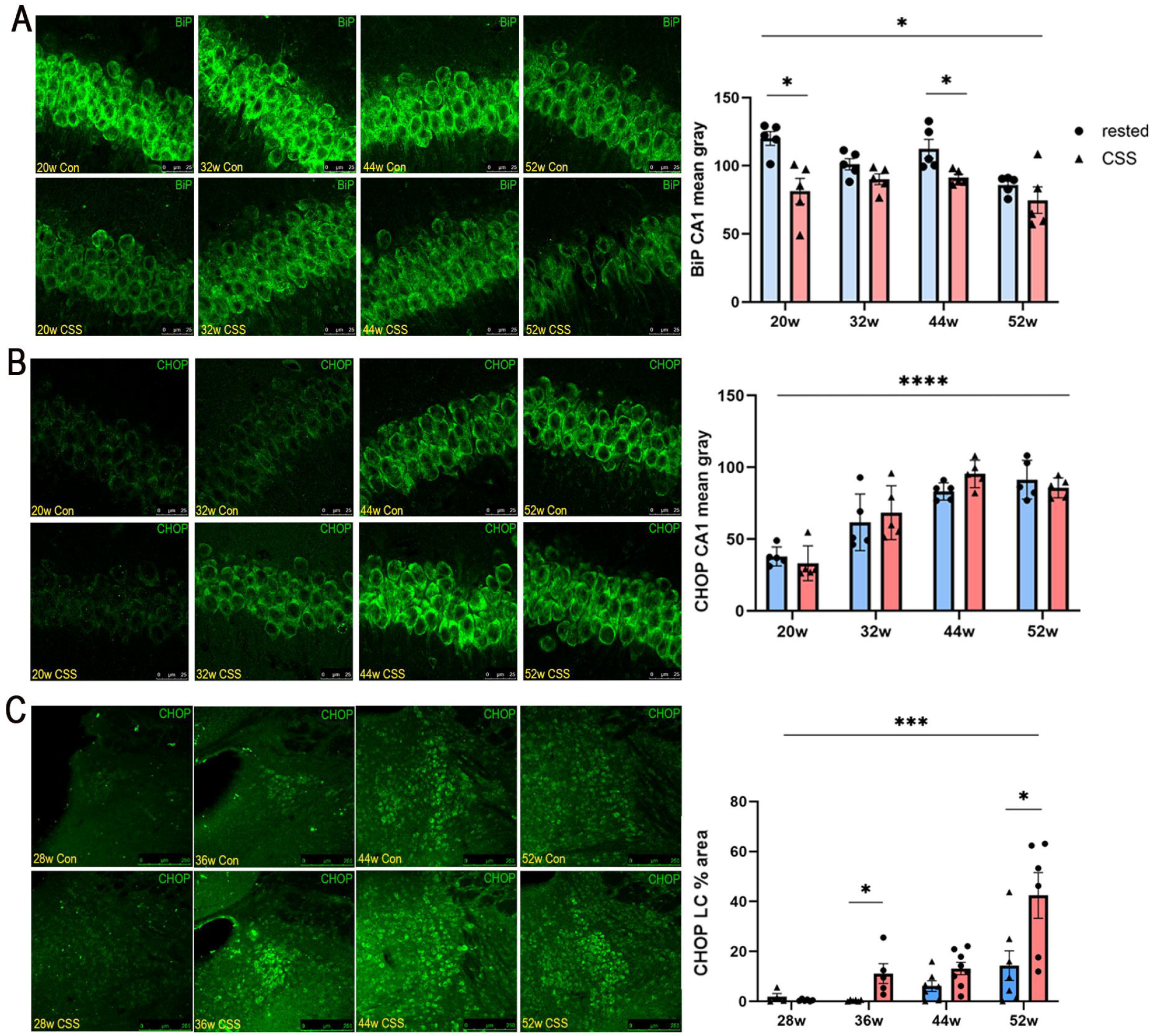
Decline in BiP and impaired proteostasis precedes cognitive impairment. Representative confocal images of the hippocampus. **A**: BiP immunofluorescent images in the CA1; **B**: CHOP immunofluorescent images in the CA1; **C**: CHOP immunofluorescent images in the LC. (Immunofluorescence data presented as mean ± SE percent area of BiP and CHOP immunofluorescence within hippocampal and LC sections; n=4-9 animals per group; two-way ANOVA for multiple comparisons, *p<0.05, ***p<0.001, ****p<0.0001).

Immunostaining also demonstrated that C/EBP Homologous Protein (CHOP) levels are elevated in the hippocampi of aged mice. CHOP, also known as Gadd153, is a transcription factor typically present at low levels under normal conditions but increases during cellular stress. While we did not find a significant increase in CHOP with CSS (p=0.29), we did find a marked increase in CHOP with age (Fig. 3B; two-way ANOVA, p<0.0001). Additionally, based on the CSS-induced molecular changes observed in the hippocampus and previously published literature that indicated that CSS led to a loss of LC neurons (Owen et al., 2021), we also examined the LC for CHOP, a notable marker of sustained ER stress. We found that CHOP was significantly increased with CSS at 36 and 52-weeks (Fig. 3C; T-test, p=0.015, p=0.022). Additionally, CHOP levels significantly increased with age in LC neurons (Fig. 3C; 2-way ANOVA, p=0.0003).

### The integrated stress response (ISR) effector, ATF4, is increased in CSS mice at 28 weeks of age

Given that we observed a divergence in cognitive behavior between the rested and CSS mice at 28 weeks of age, we used immunofluorescence to examine hippocampal sections at the 28-week timepoint for UPR and ISR markers, p-PERK, p-eIF2α, and ATF4. It is known that sustained ER and proteostatic stress leads to PERK activation, phosphorylation of eukaryotic translation initiation factor 2α (eIF2α), inhibition of global protein translation, and induction of the ISR effector, activating transcription factor 4 (ATF4). Attenuation of protein synthesis and induction of ISR are associated with impaired cognition (Bertolotti et al., 2000; Costa-Mattioli et al., 2007; Costa-Mattioli and Walter, 2020). While immunofluorescence data revealed a non-significant increase of 72. % in phosphorylated PERK in the CA1 of the CSS mice (Fig. 4A; T-test, p=0.23) and a non-significant increase of 39.8% in peIF2α levels in hippocampal tissue lysates of the CSS mice (Fig. 4C; T-test, p=0.26), ATF4 was significantly increased in the hippocampus of CSS mice (Fig. 4B; T-test, p=0.019). Given the non-significant change in p-eIF2α, we additionally assessed growth arrest and DNA damage-inducible protein 34 (GADD34), a phosphatase that dephosphorylates p-eIF2α. Notably, GADD34 levels were significantly increased in CSS mice (Fig. 4D; T-test, p = 0.02).

**Fig. 4:**
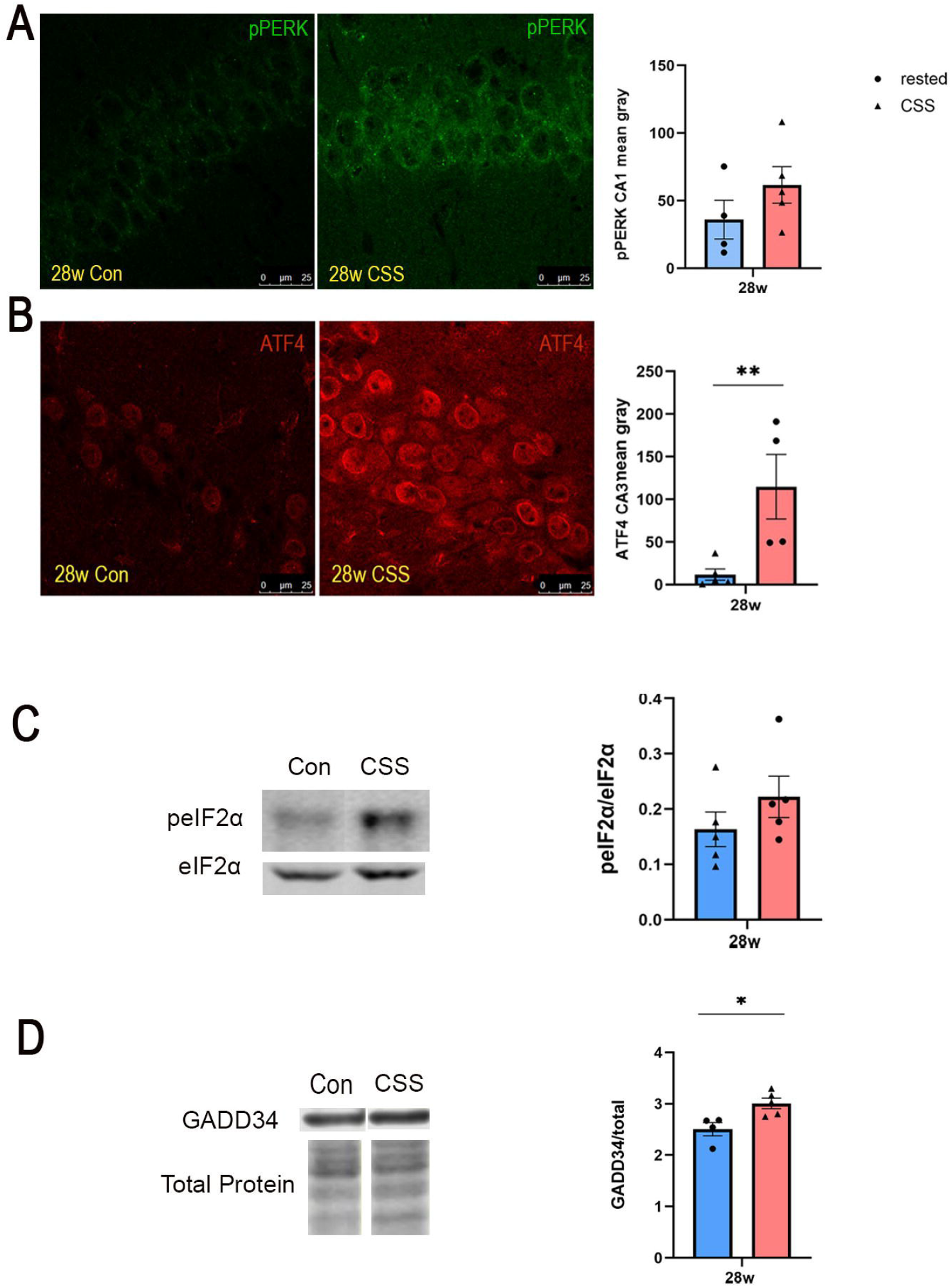
CSS leads to increased activation of notable integrated stress response (ISR) marker ATF4 at 28 weeks of age. Representative confocal images of the hippocampus across groups. **A**: PERK immunofluorescent images in the CA1; **B**: ATF4 immunofluorescent images in the CA3 (Immunofluorescence data presented as mean ± SE percent area of PERK and ATF4 immunofluorescence within hippocampal sections; n=4-9 animals per group; unpaired two-tailed t-test, **p<0.01). **C**: Representative western blot images. peIF2α ∼38 kDa. **D**: GADD34 ∼100 kDa (Membranes are cut along the MW markers following blocking to utilize the two-channel Odyssey system and varying molecular weights for the proteins of interest to probe for multiple markers within one membrane).

### CSS impacts synaptic and memory markers at 28 weeks of age

We next wanted to determine whether the CSS-induced behavioral changes seen at 28-weeks were associated with changes in synaptic plasticity and memory markers and thus measured BDNF and phosphorylated CREB (pCREB), both well-known markers of learning and synaptic plasticity (Bourtchuladze et al., 1994; Xiong et al., 2013). Immunostaining of hippocampal sections revealed a non-significant 12% decrease in pCREB levels in the dentate gyrus of the CSS mice compared to that from rested mice (Fig. 5A). We also found a modest decrease in BDNF in CSS hippocampal tissue lysate that just met the threshold for significance (Fig. 5B, T-test, p=0.05).

**Fig. 5:**
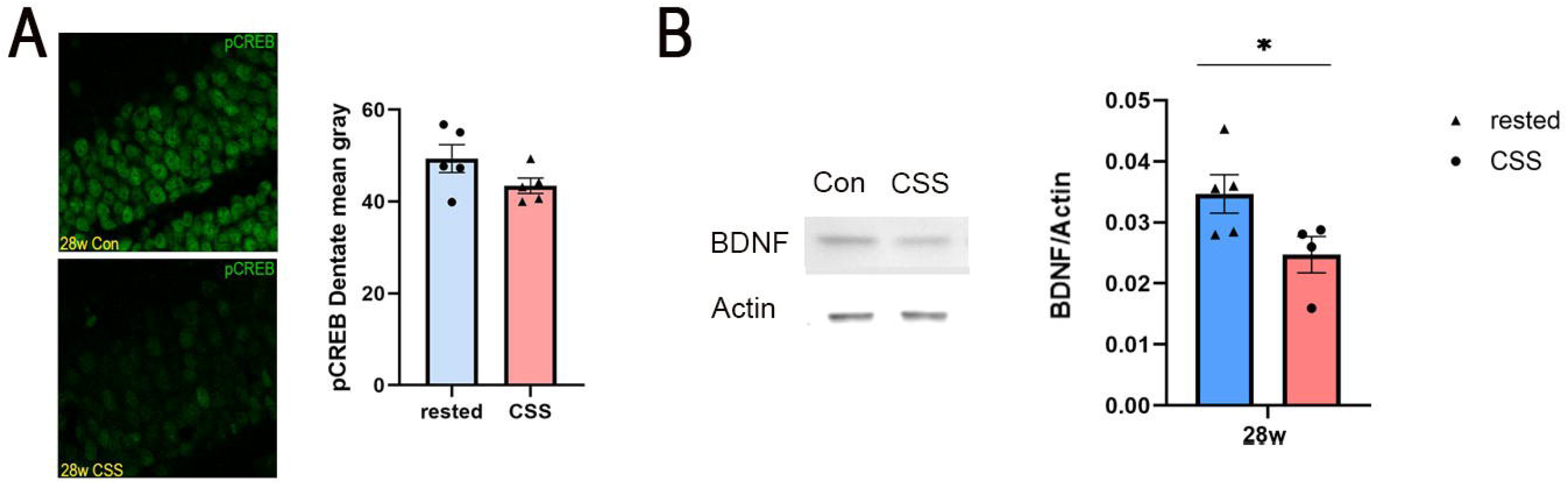
CSS leads to a decline in synaptic and memory markers at 28 weeks of age. **A**: Representative confocal images of the hippocampus. CREB immunofluorescent images in the CA1 (Immunofluorescence data presented as mean ± SE percent area of CREB immunofluorescence within hippocampal sections; n=4-9 animals per group; **B**: Representative western blot images. BDNF dimer ∼28 kDa (Membranes are cut along the MW markers following blocking to utilize the two-channel Odyssey system and varying molecular weights for the proteins of interest to probe for multiple markers within one membrane).

### Neuroinflammation increases with age

Having observed the effects CSS has on neurons, we examined other supporting structures in the brain, namely, astrocytes and microglia, at all timepoints across the study (Figures 6 and 7). Immunostaining for the astrocytic marker GFAP and microglial marker, ionized calcium binding adaptor molecule 1 (IBA1) in hippocampal sections revealed that both markers were increased with age (Fig. 6A, 6B; 2-way ANOVA, p=0.012, 0.0003). However, a count of GFAP positive astrocytes by at 28 weeks of age while higher in the CSS (21±5.8) mice than in the rested (17.4±2.2) mice, was not significant (Fig. 7A; T-test, p=0.54). Quantification of Iba1, while also increased by 36.2 % in the CSS mice sections compared to those from the rested mice at 28 weeks, was not significant (Fig. 7B; T-test, p=0.14).

**Fig. 6:**
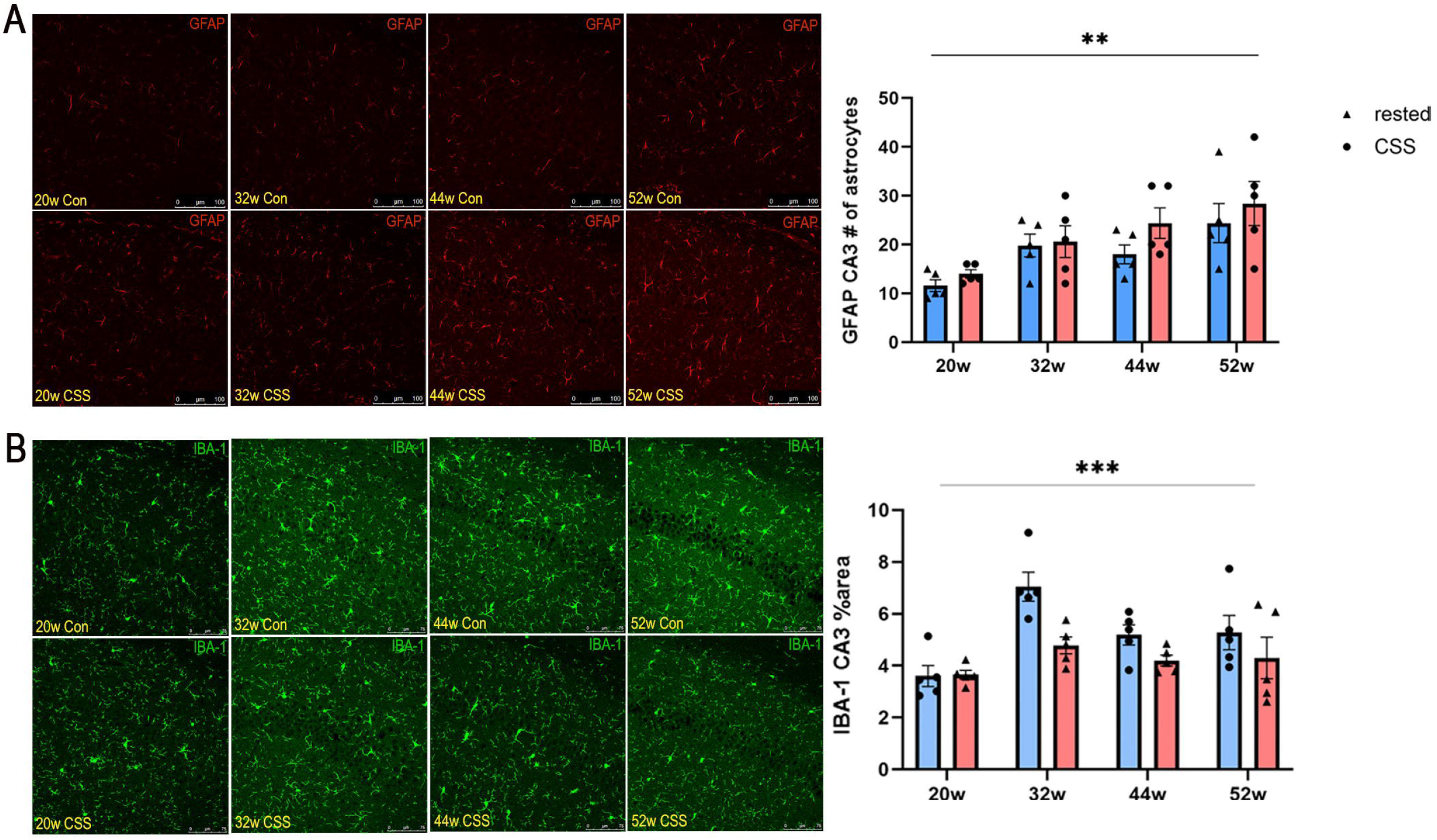
Age contributes to increased neuroinflammation. Representative confocal images of the hippocampus. **A:** GFAP immunofluorescent images in the CA3; **B**: IBA-1 immunofluorescent images in the CA3 (Immunofluorescence data presented as mean ± SE percent area of GFAP and IBA-1 immunofluorescence within hippocampal sections; n=4-9 animals per group; two-way ANOVA for multiple comparisons, **p<0.01, ***p<0.001).

**Fig. 7:**
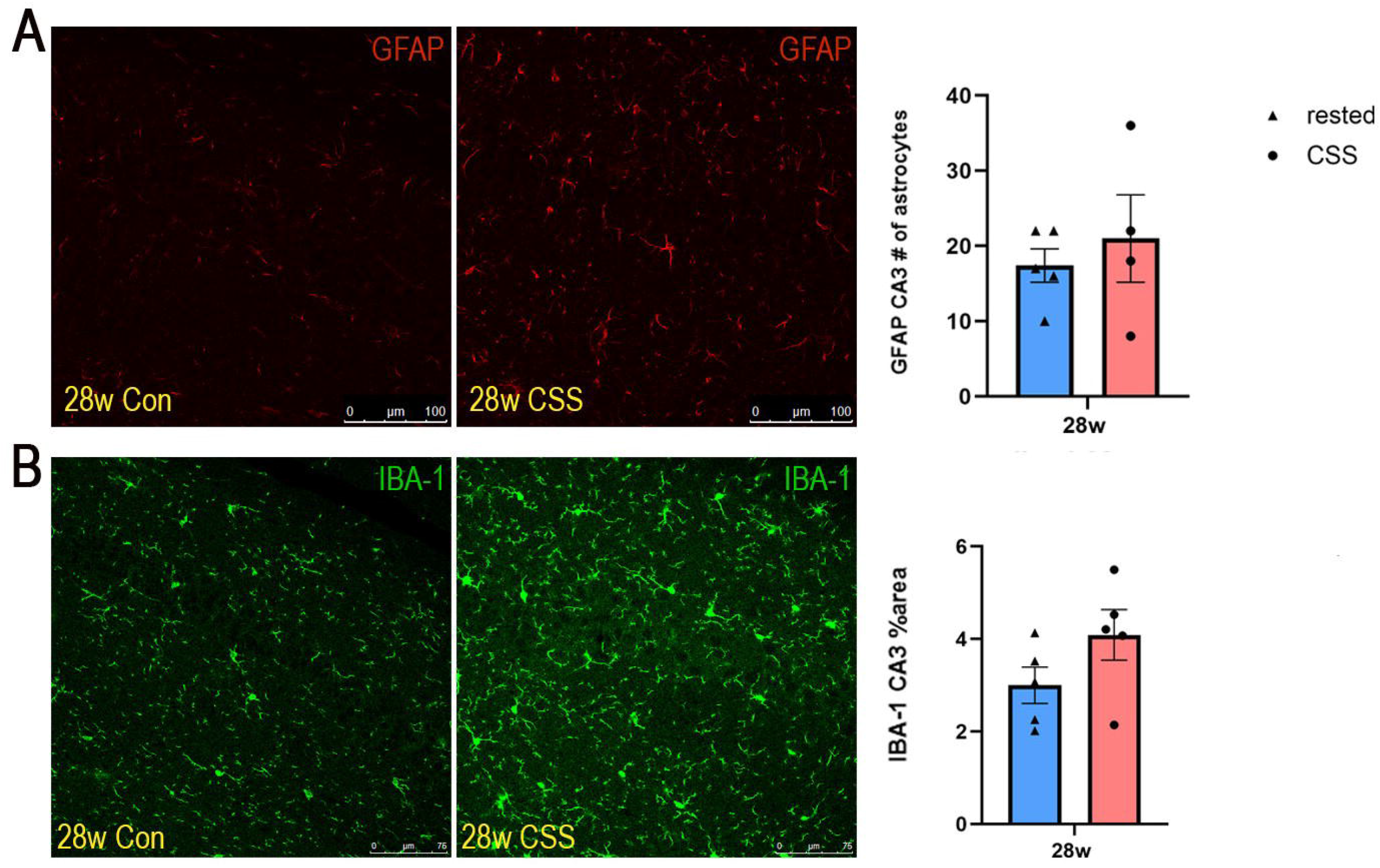
Neuroinflammation is also impacted by CSS at 28 weeks. Representative confocal images of the hippocampus. **A**: GFAP immunofluorescent images in the CA3. **B**: IBA-1 immunofluorescent images in the CA3 (Immunofluorescence data presented as mean ± SE percent area GFAP and IBA-1 immunofluorescence within hippocampal sections; n=4-9 animals per group; unpaired two-tailed t-test).

## DISCUSSION

In this study, we proposed that that chronic short sleep (CSS) induces and sustains ER stress, which leads to an integrated stress response. We also hypothesized that hippocampal memory impairments are temporally linked to this induction of ER stress. Our findings suggest that CSS does induce ER stress, which in turn affects hippocampal memory. A significant difference in spatial memory emerged at 28 weeks, when the CSS-exposed mice exhibited a decline in cognitive performance while the rested mice maintained their spatial learning abilities. Conventionally, memory formation in the hippocampus is thought to occur in three key phases: encoding, consolidation, and retrieval. Sleep plays a vital role in both encoding and consolidation by strengthening memories. This involves transforming learned information into meaningful, qualitative patterns and integrating them into existing schematic networks. Additionally, sleep facilitates generalization of these properties, detect statistical patterns, and discover underlying rules (Crowley et al., 2024). When sleep is disrupted, as with CSS, these processes are impaired, which leads to deficits in cognitive performance similar to that observed in the CSS mice.

Our findings suggest that CSS induces sustained ER stress and prolonged unfolded protein response (UPR) activation, notably preceding the cognitive decline seen at 28 weeks. Given the established role of ER stress and UPR dysregulation in memory deficits, these results suggest that such dysregulation is a key mechanism in CSS-induced cognitive decline. A key factor involved in this process is Binding Immunoglobulin Protein (BiP), a crucial chaperone and regulator of the UPR. Under regular conditions, BiP facilitates normal protein folding and prevents aggregation while simultaneously binding to transmembrane sensors that keep the UPR inactive. However, under CSS-induced ER stress, the accumulation of misfolded proteins shifts BiP binding away from these sensors, activating downstream UPR pathways. The significant decrease in BiP levels observed at 20–22 weeks serves as a critical marker of this proteostatic shift, indicating that the system is no longer able to compensate for accumulating misfolded proteins as well as leads to prolonged inhibition of protein translation though PERK activation. Because this reduction occurs immediately prior to the cognitive impairment seen in CSS mice at 28 weeks, it suggests that dysregulation of ER proteostasis is an early event that precedes and likely contributes to subsequent hippocampal dysfunction.

Given that BiP is the most upstream regulator of the UPR, its depletion likely promotes a broader maladaptive stress response through activation of the integrated stress response (ISR). While a notable increase in CHOP was not observed immediately with CSS-induced ER stress, a significant temporal increase in CHOP levels was observed with age, alongside the decrease in BiP. This suggests that while CSS-induced BiP depletion initially disrupts proteostasis, older mice are more susceptible to a sustained, maladaptive stress response that can lead to hippocampal dysfunction and eventual cell death. Although upstream ISR mediators such as phosphorylated PKR-like ER kinase (PERK) and phosphorylated eIF2α (peIF2α) showed only modest, non-significant increases, we observed a significant upregulation of the downstream activating transcription factor (ATF4). We also observed an increase in GADD34, a stress-inducible protein that recruits protein phosphatase 1 (PP1) to specifically dephosphorylate peIF2α (Novoa et al., 2001). The significant increase in GADD34 observed in CSS mice at 28 weeks likely explains the lack of significantly increased peIF2α levels at the same timepoint. The increase in ATF4 coincides with the memory deficits observed in CSS mice, indicating that chronic ISR activation may impair protein synthesis required for hippocampal memory consolidation.

Furthermore, the hippocampus is not the sole component of memory consolidation. Posterior to the hippocampus is the locus coeruleus (LC), which regulates circadian rhythm and alertness by providing norepinephrine. Importantly, LC neurons project into the hippocampus to release norepinephrine, which facilitates the encoding of significant information ((Caestecker et al., 2025). Along with the elevated CHOP levels observed in these CSS mice, and in line with prior work showing that CHOP increases in LC tissue with age and stress (Naidoo et al., 2011), CHOP levels in the LC also showed significant increases with age and with CSS at later time points (36 and 52 weeks). This suggests that the LC is highly susceptible to the maladaptive stress response associated with CSS and aging, potentially due to disruption of norepinephrine signaling to the hippocampus, further contributing to cognitive decline.

While protein synthesis halted by chronic UPR and ISR activation can impair cognition, hippocampal function is specifically degraded through the disruption of synaptic plasticity and memory-related molecular signaling. Central to these mechanisms are brain-derived neurotrophic factor (BDNF) and phosphorylated CREB (pCREB). pCREB is thought to target genes like BDNF to control memory consolidation and long-term potentiation. Previous studies have shown that mice displaying a gain of CREB function and enhanced BDNF levels exhibit enhanced memory formation (Kida, 2012). Interestingly, it has also been found that CREB expression is downregulated in AD brains (Bartolotti et al., 2016). We observed a non-significant decrease in pCREB levels and a significant decrease in BDNF levels at 28 weeks in our CSS mice. This suggests that memory is impaired not only through chronic UPR and ISR activation, but also through the direct disruption of these memory-related signaling pathways.

Beyond neuronal signaling, neuroinflammation represents another critical element of CSS’s impact on the brain. Stressors like CSS trigger the brain’s immune cells, specifically microglia and astrocytes. Microglia are seen as primary surveillance responders to homeostatic disturbance through neurotoxic and neuroprotective molecular signaling. Astrocytes are more specialized, star-shaped cells involved directly in metabolic support and blood-brain barrier regulation. When these cells transition into a reactive state, known as astrocytosis, the molecular signaling required for memory consolidation can be disrupted (Garland et al., 2022). It has also been shown that while reactive astrocytes and microglia can play a protective role by attempting to restore homeostasis, prolonged activation may contribute to synaptic dysfunction (Kim et al., 2024). Our findings indicate that as mice age, there is increased glial activation, tagged by ionized calcium binding adaptor molecule 1 (IBA1), and astrocytosis, tagged by glial fibrillary acidic protein (GFAP). While there were no significant changes at 28 weeks specifically, both exhibited a trended increase. This suggests that while reactive glia may initially play a protective role by attempting to restore homeostasis, their prolonged activation can contribute to synaptic dysfunction.

## CONCLUSION

Our results demonstrate that CSS induces proteostatic instability where the significant decrease of BiP at 20-22 weeks serves as a critical precursor to the cognitive decline observed at 28 weeks. This shift triggers a transition from an adaptive response to a maladaptive integrated stress response (ISR). While upstream initiators like pPERK and peIF2α show non-significant increases, their signals seemingly culminate into a significant upregulation of ATF4. This induction of the ISR, combined with a trended increase in neuroinflammation, blocks pCREB and BDNF signaling needed for synaptic plasticity, resulting in the significant decrease in BDNF levels observed alongside memory deficits. Aging appears to act as a cumulative stressor that prevents the resolution of ER stress, driving the significant induction of CHOP and neuroinflammation. This age-related maladaptive effect identifies a narrow therapeutic window between 20-22 and 28 weeks, where intervening in the UPR with a chaperone drug like 4-phenylbutyrate might prevent the transition from manageable cellular stress to permanent age-associated memory loss.

## MATERIALS AND METHODS

### Mice

Animal experiments were conducted in accordance with the guidelines of the University of Pennsylvania Institutional Animal Care and Use Committee. Mice were maintained as previously described (Chellappa et al., 2019; Naidoo et al., 2008). Mice were housed at 23°C on a 12:12 h light/dark cycle and had ad libitum access to food and water.

#### Chronic Short Sleep (CSS) Protocol

The CSS protocol was carried out at the University of Pennsylvania in accordance with the National Institutes of Health Office of Laboratory Animal Welfare Policy and the University of Pennsylvania’s Institutional Animal Care and Use Committee as described in (Owen et al., 2021)with a few modifications. Briefly, 8-week-old male C57BL/6J (Jackson Laboratory) mice maintained in groups of 5 (Total n=80) were subjected to CSS or rest control conditions for a period of 8 weeks. CSS was performed by placing mice and their littermates in novel enriched environments, where climbing toys were exchanged whenever a mouse in the paradigm became quiescent (Gompf et al., 2010). During CSS, mice were continuously observed by one or two researchers, each watching mice directly for 1–2 h shifts to ensure eight continuous hours of active wakefulness in each mouse. This CSS paradigm has been shown to not elevate blood corticosterone levels (Zhang et al., 2014). Mice were subjected to CSS for 3 consecutive days of each week for the first 8 h of the lights-on period, when mice typically sleep (see Figure 1). Rested controls were exposed to the same environment for 1 hour per day for 3 days per week for 8 weeks.

### Spatial Objection Recognition (SOR) Test

The Spatial Objection Recognition test is well-established hippocampal-dependent spatial memory test (Bevins and Besheer, 2006; Cavoy and Delacour, 1993; Hafycz et al., 2022). The mice are placed together for an hour in the testing apparatus for three consecutive days prior to testing to acclimate to the container. Testing occurs in two phases. The first is the training phase where two identical objects are placed on one side of the apparatus. Mice are placed individually in the apparatus for 10 minutes and all interactions (smelling, touching, etc.) are counted. After training, the mice are returned to their home cage and left undisturbed for 90 minutes. Before the start of the testing phase, one of the identical objects is moved to the opposite side of the apparatus. Mice are placed in the apparatus during the testing phase similar to the training phase, but only for 3 minutes. Interactions with each object are measured again. Discrimination index calculations were performed as previously described as a measure for how well the mice distinguish between the moved and unmoved object (Sivakumaran et al., 2018).

### Immunohistochemical Assays

Post-fixed half-brain coronal sections were sliced at 40 μm using a cryostat as previously described (Hafycz et al., 2022; Zhu et al., 2007). Every other section was placed in 24-well plates containing cryoprotectant for free floating immunohistochemistry staining and stored at -20ºC, as previously described (Hafycz et al., 2022; Naidoo et al., 2018). For all markers, we compared n=4-5 in each of the groups.

### Immunofluorescence (IF)

Immunofluorescence staining was conducted as previously described (Hafycz et al., 2022; Naidoo et al., 2011). Primary antibodies are as follows: pCREB (ser133) (1:300, Cell Signaling 87G3); CREB (1:200, Cell Signaling 86B10); pPERK (Thr980) (1:200, Bioss bs-3330R); ATF4 (1:500, ProteinTech, 60035-1-Ig); BiP/anti-KDEL (1:1000, Enzo Life Sciences ADI-SPA-827F); GFAP (1:300, Cell Signaling 3670); IBA-1 (1:300, Cell Signaling 17198); Tyrosine Hydroxylase (1:1000, Abcam, ab113). Secondary antibodies are as follows: Alexa Fluor 488 donkey anti-rabbit IgG (1:500); Alex Fluor 594 donkey anti-mouse IgG (1:500); Alexa Fluor 488 donkey anti-mouse IgG (1:500); Alexa Fluor 594 594 donkey anti-Sheep IgG (1:500).

### Quantitative Analysis of IF Images

Confocal images were acquired as previously described (Hafycz et al., 2022; Owen et al., 2021), using Leica SP5/AOBS microscope. Confocal laser intensities, nm range, detector gain, exposure time, amplifier offset, and depth of the focal plane within sections per antigen target were standardized across compared sections. Confocal images were quantified as previously described (Hafycz et al., 2022). Briefly, 3-4 sections were imaged per animal (n=4-5 animals per group). Using ImageJ software, the images were converted to an 8-bit grayscale with detection threshold standardized across all images to detect percent areas. The percentage area and mean gray covered within the target region were measured and the averages of these measures for each mouse were analyzed.

### Western Blot

Frozen brain tissue was prepared for western blot assays as previously described (Naidoo et al., 2008; Naidoo et al., 2018). Briefly, brain tissue was homogenized on ice with lysis buffer containing protease inhibitors. After centrifugation, protein concentration for each sample was determined with a BCA protein assay and samples were prepared such that each contained 20µg of protein. SDS-PAGE gels were run as previously described (Naidoo et al., 2008), and protein bands were imaged and quantified via infrared imaging on an Odyssey scanner (LiCor). For all markers we compared n=5-8 for each of the groups. Primary antibodies are as follows: GADD34 (1:500, Protein Tech 10449-1-AP); peIF2α (1:100, Cell Signaling 3597S); BDNF (1:500, Abcam ab108319); β-Actin (1:2000, Santa Cruz sc-47778). Secondary antibodies are as follows: LiCor IRDye 680RD Goat anti-Mouse (1:10,000); LiCor IRDye 800RD Goat anti-Mouse (1:10,000); LiCor IRDye 800RD Goat anti-Rabbit (1:10,000); Odyssey IRDye 680 Goat anti-Rabbit (1:10,000).

## Funding

R01 AG064231 (Cellular and Molecular Basis of Sleep Loss Neural Injury in Alzheimer Disease)

## Author contributions

RK performed experiments and behavioral data analyses. KS wrote the manuscript, conducted IHC, western blot experiments, and data analyses. ES conducted western blot experiments. NN conceptualized and designed the study, provided the resources, interpreted data, edited, and revised the manuscript.

## Competing interests

The authors have no conflicts of interest to declare.

## Data and materials availability

The raw data supporting the conclusions in this manuscript will be made available upon request.

## Notes

### Competing Interest Statement

The authors have declared no competing interest.

